# Terrigenous inputs link nutrient dynamics to microbial communities in a tropical lagoon

**DOI:** 10.1101/2025.06.11.659169

**Authors:** Christian John, Nyssa J. Silbiger, Thomas C. Adam, Danielle M. Barnas, Kalia S. Bistolas, Robert C. Carpenter, Noe Castañeda, Megan J. Donahue, Mary K. Donovan, Lauren N. Enright, Hannah E. Epstein, Jordan P. Gallagher, Hendrikje Jorissen, Jamie R. Kerlin, Savanah L. Leidholt, Rowan H. McLachlan, Nury Molina, Catherine A. Mullenmeister, Kyle Neumann, Julianna J. Renzi, Denise P. Silva, Kelly E. Speare, Sean Swift, Alex D. Vompe, Linda Wegley Kelly, Maya Zeff, Craig E. Nelson, Rebecca Vega Thurber, Deron E. Burkepile

## Abstract

Nutrient availability drives community structure and ecosystem processes, especially in tropical lagoons that are typically oligotrophic but often receive allochthonous inputs from land. Terrestrially-derived nutrients are introduced to tropical lagoons by surface runoff and submarine groundwater discharge, which are influenced by seasonal precipitation. Lagoon habitats are distributed along an onshore-offshore gradient; terrigenous inputs presumably diminish along the same continuum. We characterized nutrient enrichment in the lagoons of a tropical high island, Moorea, French Polynesia, using spatially distributed measurements of nitrogen content in the tissues of a widespread macroalga during the rainy season over four years. We used synoptic water column sampling to identify associations among macroalgal nutrient content and the composition of inorganic macronutrients, dissolved organic matter, and microbial communities. We paired these data with quantifications of land use in nearby watersheds to uncover links between terrestrial factors, aquatic chemistry, and microbial communities. Algal N content was highest near shore and near large, human-impacted watersheds, and lower at offshore sites. Sites with high algal N had water columns with high nitrite + nitrate, silicate, and increased humic organic matter (based on a fluorescence humification index), especially following rain. Microbial communities were differentiated among nearshore habitats and covaried with algal N and water chemistry, supporting the hypothesis that terrigenous nutrient enrichment shapes microbial dynamics in otherwise oligotrophic tropical lagoons. This study reveals that land-sea connections create nutrient subsidies that are important for lagoon biogeochemistry and microbiology, indicating that changes in land use or precipitation will modify ecosystem processes in tropical lagoons.

## Introduction

Nutrients regulate primary production, trophic interactions, and decomposition (Tilman et al. 1982; Cleveland et al. 2006; Moore et al. 2013). Tropical lagoon ecosystems are relatively oligotrophic (Johannes et al. 1972), but they are among the most productive ecosystems on the planet in spite of their relatively low nutrient concentrations (D’Elia and Weibe 1990; Atkinson and Falter 2003). Yet, terrestrial factors such as the presence of nitrogen-fixing vegetation and human modification of the landscape enrich nutrients in many lagoons worldwide (Carlson et al. 2019). For such oligotrophic systems, terrigenous nutrient subsidies alter baselines of limiting resources and affect community dynamics (Dong et al. 2017; Adam et al. 2021). Microbes process a large proportion of organic material in aquatic systems (Azam et al. 1983), but are highly sensitive to nutrient dynamics (Henson et al. 2018), and therefore represent an important fulcrum by which nutrients impact ecosystem function.

Water column nutrient concentrations in near-shore ecosystems can fluctuate over short timescales due to ephemeral variation in precipitation runoff, submarine groundwater discharge, and ocean currents (Hench et al. 2008; Strauch et al. 2015; Santos et al. 2021; Silbiger et al. 2025). The delivery of terrigenous nutrients to nearshore ecosystems, especially those from anthropogenic sources, is often episodic, with increased flux generally coinciding with high rainfall events (Fong et al. 2020; Trefry and Fox 2021). Thus, highly dynamic water column chemistry may mask long-term nutrient patterns due to short-term environmental variability. Furthermore, because nutrient flux is important and often limiting for organisms, water column nutrients are subject to rapid biological uptake and therefore may not be indicative of nutrient availability. Because water column nutrients are integrated by autotrophs (e.g. Hein et al., 1995), nutrient concentrations in the tissues of primary producers can be useful for assessing patterns of nutrient availability over longer time scales (Fong et al. 1994; Adam et al. 2021). Together, synoptic sampling of the water column and bioindicators (such as producer tissues) can be a powerful way to integrate complementary perspectives of nutrient patterns across different temporal and spatial scales (Salo and Salovius-Laurén 2022).

Microbes in coastal oceans contribute to important ecosystem functions by fixing nutrients and cycling organic matter (Gupta et al. 2017). Water column microbial communities are central to biogeochemical cycling of lagoon systems and are important components of the trophic structure of nearshore environments (Nelson et al. 2023). Microbial communities can be shaped by patterns of nutrient availability, especially in oligotrophic systems (Koyama et al. 2014; Montiel-González et al. 2017; Haber et al. 2022). Therefore, water column microbes may be sensitive to anthropogenic factors in typically nutrient-poor coastal ecosystems if humans upend historic patterns of water chemistry, such as in cases of eutrophication (Aldeguer-Riquelme et al. 2022; Deng et al. 2024). The spatial arrangement of nutrient inputs is thus a potentially important driver of microbial community dynamics, and by extension, ecosystem function in tropical lagoons.

One important axis of physicochemical variation in tropical high island lagoons lies along the spatial gradient from shore to reef crest (Shuler et al. 2019). The balance of terrestrial and oceanic inputs varies across this onshore-offshore continuum (Leng et al. 2023). For example, terrigenous nutrients may be found at higher concentrations near shore and lower concentrations far from shore due to biological uptake, settling, and dilution processes (Lapointe and Clark 1992). Discrete reef habitat types (including fringing reef, back reef, and reef crest) are arranged within the onshore-offshore gradient, so reef-associated microbes are subject to nutrient dynamics associated with habitat location (Wang et al. 2019). Complicating this relationship, terrestrial features such as catchment size and land use may drive patterns of variation in nutrient inputs perpendicular to the onshore-offshore gradient due to their impact on the chemistry of water run-off (Knee et al. 2010). Thus, microbial communities in similar habitat types may face different environmental pressures if they are adjacent to or downstream of different terrestrial features.

Here we examined spatiotemporal relationships among macroalgal tissue chemistry, water column chemistry, and water column microbial communities from the lagoons of Moorea, French Polynesia, a tropical high island in the South Pacific. We created a multiannual dataset of algal tissue nitrogen content, water column dissolved inorganic and organic nutrients, and planktonic microbial community composition at nearly 200 sites across the lagoon spanning the habitat continuum from fringing reef to reef crest. We asked the following questions: 1) What is the spatiotemporal structure of lagoon nutrient availability? 2) Does water chemistry in nearshore lagoon environments derive from terrestrial sources, and is it associated with land use? And 3) To what extent are lagoon microbial communities influenced by the availability of nutrients? We hypothesized that nutrient patterns in both algal tissues and water would be associated with proximity to shore and patterns of land use, that precipitation would increase lagoon nutrients, and that microbial communities would differ by habitat in association with patterns of nutrient availability.

## Materials and Methods

### Site description and sampling overview

In this study, we examined patterns of nutrient availability across the seascape and their potential impacts on water column microbiomes in the lagoons of Moorea, French Polynesia (17.5°S, 149.8°W). Research was conducted at the University of California, Gump Research Station on the north shore of Moorea. Moorea is a steep, volcanic island; its ∼17,000 inhabitants reside mostly within 0.5km of shore. The limited low-elevation terrain further inland is used, in large part, for agriculture: over 1/3 of land below 50m elevation and more than 500m from shore on Moorea is agricultural. The water surrounding Moorea is oligotrophic, and its lagoons are protected from the open ocean by a barrier reef 1-2km from shore that is broken by several reef passes, often at the mouths of bays (Fig. 1) (Adam et al. 2021). Lagoon habitats consist of shallow fringing reefs that ring the island, often adjacent to deepwater channels. Offshore of these deepwater channels are backreef habitats ∼1-3m deep that consist of a matrix of sand and coral patch reefs that extend to the reef crest.

**Figure 1.**
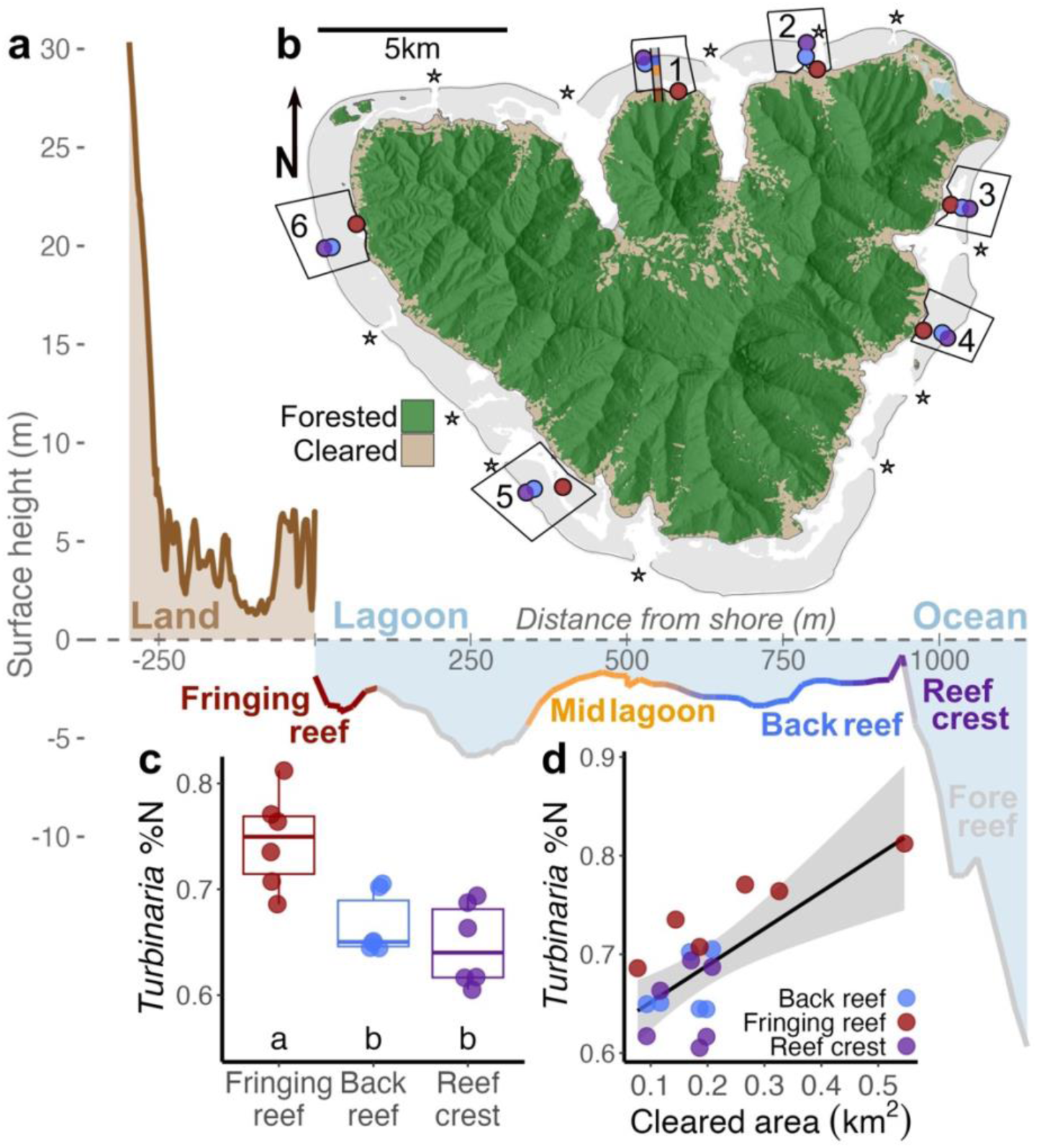
Shallow lagoons encircle Moorea, extending out to the reef crest beyond which bathymetry drops precipitously on the fore reef. A cross-sectional lagoon depth contour illustrating onshore-offshore habitat structure (a) was drawn from the transect at MCR LTER site 1, shown in map (b) with a shaded rectangle. Reef passes are indicated with stars in (b). Tissues of the brown alga *Turbinaria ornata* sampled from three reef habitats at six sites around Moorea (2007-2022) had higher %N at fringing reef sites than back reef or reef crest sites (c; annotated letters show significant differences [p<0.05] among habitat after accounting for differences among years and sites using estimated marginal means). *Turbinaria* tissue %N was higher at sites near watersheds with more land clearing (d).

A 16-year time series (2007–2022) from the Moorea Coral Reef Long-Term Ecological Research (MCR LTER) platform demonstrates that tissue nitrogen content of the brown macroalga *Turbinaria ornata* (hereafter, “*Turbinaria*”) has been consistently higher on fringing reefs than on areas of the lagoon farther offshore (Fig. 1c,d; Moorea Coral Reef LTER and Carpenter 2023).

However, these measurements are necessarily limited to a relatively small number of sites (3 habitats within each of 6 sites) around the island, and their deliberately homogenous arrangement relative to fine-scale spatial gradients near and far from terrigenous nutrient sources prevent analysis of heterogenous local terrigenous influences. Thus, we designed a fine-scale sampling campaign to complement the long-term time series, sampling tissues of *Turbinaria* across 204 sites separated by 349±144m (mean±s.d.) around the lagoon 4 times across a 7-year period (2016, 2021, 2022, and 2023). In the long-term dataset, *Turbinaria* samples were collected from fringing reef, backreef, and reef crest sites, and in the fine-scale dataset, samples were collected from fringing reef, mid-lagoon, backreef, bay, and reef pass sites. We used the elemental weight percent nitrogen in *Turbinaria* tissue (“*Turbinaria* %N”) as a proxy for nutrient availability to ask questions about patterns of nitrogen enrichment across the seascape of Moorea.

Additionally, in 3 of the 4 years (2021, 2022, and 2023), we quantified water column chemistry (nitrite + nitrate, phosphorus, silicate, and fluorescent dissolved organic matter [fDOM] components and indices) and water microbial communities (via 16s rRNA sequencing) from water samples taken synoptically with *Turbinaria* samples. We compiled archived meteorological data to summarize daily precipitation across the duration of the fine-scale *Turbinaria* sampling campaign. Finally, we combined fine-scale topography and land cover datasets to understand land use variability across Moorea’s watersheds. We used these additional data to determine how time-integrated measurements of nitrogen content in *Turbinaria* tissues related to instantaneous measurements of water column nutrients, and detect links among land use, precipitation, lagoon nutrients, and microbial communities.

### Field collections and sampling processing

#### *Turbinaria* sampling and processing

To characterize nutrient dynamics, we used nitrogen content of tissues sampled from *Turbinaria* around the lagoons of Moorea (*Turbinaria* %N). A long-term, but spatially limited dataset (Moorea Coral Reef LTER and Carpenter 2023) was used to contextualize nutrient dynamics over a 16 year history (2007–2022). Sampling for this long-term dataset included fringing reef, back reef, and reef crest habitats, and occurred during May in each year. A finer-scale, island-wide sampling occurred over approximately three weeks during each of four recent sampling periods (May 2016, May 2021, April 2022, and April 2023); due to logistical constraints, the number of sites varied from 170-198 in each sampling period (see Supplementary materials S1 for sampling scheme). The sampling design included fringing reef, mid-lagoon, back reef, reef pass, and bay sites. *Turbinaria* stores surplus N in its tissues and, consequently, has been used previously to index N availability over weeks to months in this system (Donovan et al. 2020; Adam et al. 2021). For the long-term time series, thalli from 5-10 different patches of *Turbinaria* were collected at each site during each sampling period. For the lagoon-wide dataset, thalli from 10 patches were collected at each site during every sampling period. In both sampling frameworks, one blade was sampled 5 cm from the apical tip for each thallus from each site. Each blade was cleaned of epiphytes and rinsed with fresh water, then dried at 60°C, and ground to a fine powder. Total N content was determined via elemental analysis using a CHN elemental analyzer (NA1500, Carlo-Erba) at the University of Georgia, Center for Applied Isotope Studies for fine-scale *Turbinaria* samples collected in 2016, and at the University of California, Santa Barbara (UCSB), Marine Science Institute Analytical Laboratory for the long-term *Turbinaria* time series and for the fine-scale *Turbinaria* samples collected in 2021, 2022 and 2023 (CEC 440HA, Exeter Analytical). In 2016, 20 samples were sent to both labs for cross-correlation, showing that %N from the two labs were highly positively correlated (see Supplementary materials S2).

#### Water column sampling

At each fine-scale *Turbinaria* sampling site, ∼1.0L of seawater was collected by snorkelers, 1m off the benthos, using a thrice-rinsed 1.5L low density polyethylene bag (Whirl-Pak; Nasco Sampling LLC, Wisconsin, USA). Samples were transported to a nearby boat and filtered using a 60mL syringe and a sterile 0.2um polyethersulfone Sterivex filter unit (Sterivex; model SVGP01050 Millepore Sigma). For water column nutrients, 15ml of the filtrate was collected into a sterile polypropylene vial. For fDOM, 20ml of the filtrate was collected into an amber glass vial. A total of 800 ml of seawater was filtered through the Sterivex filter for eDNA. Sterivex were then fully dried by forcing air through the filter to expel any remaining liquid. The dried Sterivex filter was then stored in a sterile, pre-labelled Whirl-Pak, and both it and the water sample vials were placed on ice in a cooler during transport back to the Gump Research Station. The Sterivex filters and nutrient samples were then frozen at -40°C, and the fDOM samples were stored at 4°C. All Sterivex filters were transported frozen to the Department of Microbiology at Oregon State University (OSU), all nutrient samples were transported frozen to UCSB, and all fDOM samples were shipped and stored refrigerated until processing at the University of Hawaiʻi.

#### Water column chemistry processing

Analyses of water column nutrients (nitrite + nitrate, phosphate, and silicate) were conducted using flow injection with an automated ion analyzer (QuickChem 8500, Zellweger Analytics) at the UCSB Marine Science Institute Analytical Laboratory. Analyses of water column fDOM were conducted using three-dimensional excitation-emission scanning fluorometry following (Nelson et al. 2015). A set of fDOM components (organic compounds) and fDOM indices (derived values from combinations of fDOM components) were selected to describe lagoon organic chemistry dynamics (summarized in Nelson et al. 2015). fDOM components included visible, ultraviolet, and marine humic-like compounds (large complex molecules created through decomposition of organic matter); and lignin-like, tryptophan-like, and tyrosine-like compounds (smaller molecules associated with plant matter in the case of lignin and proteinaceous material for the others). fDOM indices included HIX (Humification Index, a measure of the total amount of humics in the sample), BIX (Biological Index, inversely associated with the age of organic compounds), M:C (the ratio between marine humic-like and visible humic-like compounds, associated with marine sources), and FI (Fluorescence Index, inversely associated with terrestrial sources (Nelson 2009)). For more details on water chemistry sample processing, see Supplementary materials S3.

#### Water column microbial community processing

Analyses of the taxonomic profiles of water column microbial communities (archaea and bacteria) were conducted using 16S rRNA amplicon sequencing. DNA was extracted using a modified DNeasy Blood and Tissue Kit protocol (Qiagen, North Rhine-Westphalia, Germany). Polymerase Chain Reaction (PCR) was performed to amplify the V4 region of the 16S rRNA gene using the primer pair 515F (5′-GTGYCAGCMGCCGCGGTAA-3′) (Parada et al. 2016) and 806R (5′-GGACTACNVGGGTWTCTAAT-3′) (Apprill et al. 2015). Primers were synthesized to incorporate both Illumina adapter sequences and unique dual indices (barcodes), following the one-step PCR protocol following the Earth Microbiome Project (Caporaso et al. 2012). Libraries were pooled at equimolar concentrations and sequenced paired-end 2×250 bp sequencing using the Illumina MiSeq system at Oregon State University’s Center for Qualitative Life Science (CQLS).

For additional details on DNA extraction and amplification, see Supplementary materials S4 Demultiplexing of sequence data was performed by CQLS. Sequencing data were processed using QIIME2 v.2023.7 (Bolyen et al. 2019). Primers were removed using the cutadapt plugin (Martin 2011). Quality control of reads, including dereplication, denoising, chimera removal, trimming, and the identification of amplicon sequence variants (ASVs) were done using the DADA2 plugin (Callahan et al. 2016). The taxonomic assignment of the ASVs was conducted using the classify-sklearn plugin with a Naive Bayes classifier (Bokulich et al. 2018) trained on the SILVA V.138.1-99% OTUs reference database (Quast et al. 2013), specific to the 515F–806R region. A rooted phylogenetic tree was constructed using the align-to-tree-mafft-fasttree pipeline within QIIME 2. The output files, including the taxonomy, representative sequences, and ASV table, were converted into a phyloseq object using phyloseq v.1.44.0 (McMurdie and Holmes 2013). Sequences lacking assignment beyond the kingdom level, or classified as Chloroplasts, Eukaryota, or unassigned kingdom, were removed prior to analysis. To minimize sequencing noise, singleton ASVs were excluded from subsequent analyses. Potential negative control contaminants of sequencing data were removed as well using the R package decontam (Davis et al. 2018) based on the prevalence method with a threshold of 0.5; in total, less than four contaminants were removed. Samples were rarefied to 10,500 reads per sample to standardize sequencing depth and enable comparative analyses of community structure.

#### Environmental data

Geospatial parameters, patterns of land cover, and precipitation were summarized from publicly available data sources. For each of the lagoon sampling locations, a relevant watershed on Moorea was delineated in order to characterize potential terrestrial stressors unique to each lagoon site. The watershed delineation scheme for each sampling site is as follows: (1) the nearest point on shore was identified and distance between the sampling location and the nearest shore point was calculated, (2) a local shoreline was isolated using a 500m radius around the nearest shore point, and (3) shorelines were determined using cadastral data from the Direction des affaires foncières (DAF) de la Polynésie française vector map database (<https://www.data.gouv.fr/fr/datasets/base-de-donnees-cartographique-vectorielle/>). The watershed for the local shoreline was then delineated using a fine-scale terrain dataset (Gruen et al. 2017) with WhiteboxTools v2.4.0 (Lindsay 2016; Wu and Brown 2024). Land cover within each watershed was summarized from a recent fine-scale land cover dataset (1.4m-pixels, Neumann et al 2025; <https://portal-s.edirepository.org/nis/metadataviewer?packageid=knb-lter-mcr.6005.3>). Land cover was binarized into the classes “cleared” (for development, agriculture, etc.) and “forested”, which minimized errors in the classification scheme (Neumann et al. 2025). The total surface area of pixels classified as “cleared” (km^2^) was used to summarize land use intensity within each watershed. Precipitation data were drawn from the 6 meteorological stations that were all operational on Moorea during the sampling period (2016– 2023) via <data.gouv.fr>. Daily precipitation (mm) was summarized at the island-wide scale using mean daily precipitation across all 6 stations.

### Statistical analyses

#### Macroalgal nutrient enrichment analysis

The 16-year time series of *Turbinaria* sampling was used to contextualize patterns of nutrient enrichment over decadal timescales in Moorea. A linear mixed effects model was constructed with *Turbinaria* %N as the response variable, habitat as the predictor, and site identity and year as random intercepts. Differences among habitat were determined using estimated marginal means. To test for associations among cleared area in upstream watersheds and lagoon nutrient enrichment, a linear model was fit with overall mean *Turbinaria* %N as the response variable and distance from shore and cleared land as predictors.

The fine-scale, island-wide sampling campaign was used to characterize spatial variation in lagoon nutrient enrichment around Moorea. In an effort to maximize the number of sites that could be classified according to their nutrient dynamics, missing values (due to weather or site inaccessibility) were imputed based on island-wide sampling and site information. First, site-wise *Turbinaria* %N was modeled over the entire dataset as a function of site ID and island-wide mean *Turbinaria* %N during that sampling period in a linear model. Then, island-wide mean %N for each sampling period was recalculated taking into account the coefficient estimates of sites that were missing during that period. Finally, missing site-wise *Turbinaria* %N values were estimated with model predictions using site ID and the recalculated island-wide mean %N as predictors. Missing-value imputation had a negligible effect on the results of the study (Supplementary materials S5).

Individual site-year *Turbinaria* %N measurements were used as an instantaneous measure of nutrient enrichment, and mean *Turbinaria* %N for each site over the full time series was used to summarize long-term nutrient dynamics. Differences in instantaneous *Turbinaria* %N measurements among reef habitat types (i.e., fringing, mid-lagoon, and back reefs, as well as bays and reef passes) were assessed using a linear mixed effects model with year as a random intercept then estimating marginal means of habitat groups. In our dataset, mean *Turbinaria* %N decreases with increasing distance from shore, so residuals from this relationship were taken to assess which sites were enriched or depleted in nutrients over long timescales. To visualize variation in nutrient enrichment around the lagoons, residual *Turbinaria* %N values were spatially interpolated using kriging in a spherical variogram model with a resolution of 1000 pixels (corresponding to pixels with dimensions ∼17×20m) using the R package ‘gstat’ (Gräler et al. 2016). Assumptions for the spherical variogram model (Supplementary materials S6) – stationarity and isotropy – were evaluated based on model convergence and by checking for consistent variogram fit across the breadth of sample distances, and for consistent variogram structure among all cardinal and intercardinal directions (i.e. 0°, 45°, 90°, and 135°).

#### Water column chemistry analysis

All dissolved nutrient parameters (nitrite + nitrate, silicate, and phosphate) were log transformed prior to statistical analyses. Two water column nitrite + nitrate samples were extreme outliers in 2022; both were over 7 standard deviations above the mean and were removed from the analyses. Differences in instantaneous water column N among reef habitat types were assessed using a linear mixed effects model with year as a random intercept then estimating marginal means of habitat groups. To investigate whether spatial variability in water column N predicted that of *Turbinaria* %N, these variables were compared in a linear model with *Turbinaria* %N as the response variable and water column N, Year, and their interaction as predictors. Slope homogeneity among years was investigated in an analysis of covariance (ANCOVA) using the R package ‘rstatix’ (Kassambara 2023). To test whether differences in water column nutrients and fDOM among habitats were consistent among years, linear models were used where the water column nutrient or fDOM parameter was the response variable, and habitat type, year, and their interaction were used as predictors; interannual consistency was interpreted as a nonsignificant fixed interaction effect.

A principal components analysis of log-transformed and standardized (z-scored) water column nutrients, fDOM components, and fDOM indices were used to derive two primary axes of variation for water column chemistry. Differences in water column chemistry among habitat types were investigated using PERMANOVA with Euclidean distance of water chemistry compositional differences among samples as the response variable. We considered log-transformed distance from shore (m), log-transformed area cleared of forest in the watershed (km^2^), log-transformed mean *Turbinaria* %N, and detrended *Turbinaria* %N (after accounting for distance from shore) as environmental variables. Significant associations were visualized in the ordination space using vectors drawn from envfit() (Oksanen et al. 2025).

#### Effects of recent precipitation on lagoon nutrients

Effects of terrestrial precipitation on lagoon chemistry are likely lagged due to the latency of runoff from land to sea. To identify the characteristic timescale of precipitation effects on lagoon nutrients, cumulative precipitation was calculated over a window between 1 and 60 days leading up to each sampling event (i.e., cumulative rain the day of sampling, cumulative rain the day before and the day of sampling, etc., up to cumulative rain over the two months prior to sampling). Two linear mixed effects models were constructed for each of the 60 different time windows, one with *Turbinaria* %N as the response and the other with water column nitrite + nitrate as the response. For both models, cleared land area and cumulative precipitation were used as predictors, and site was used as a random effect. Then, short (3 days) and long (30 days) timeframes were selected to discern immediate and direct, vs. chronic and cumulative, effects of terrestrial inputs into lagoons. Contrast plots were used to examine impacts of land clearing on lagoon nutrients in the mixed effects models for these two timeframes.

#### Microbial community analysis

Water column microbial community data resolved as amplicon sequence variants were aggregated to the genus level after classification to facilitate comparisons across years. A Bray-Curtis distance matrix was calculated for the microbial communities and ordinated using principal coordinates analysis (PCoA) a form of multidimensional scaling (MDS). Differences in microbial communities among habitat types were determined using PERMANOVA implemented with adonis2() and ordination correlations with environmental variables were visualized using envfit() in the ‘vegan’ package (Oksanen et al. 2025). Differences in microbial community dispersion among habitat types were determined using PERMDISP2 implemented with betadisper(). The most parsimonious model explaining differences between microbial communities was identified using AICc model selection framework after removing highly correlated sets of environmental variables (using a variance inflation factor cutoff of 5) in the ‘AICcPermanovà package (Corcoran 2023). Candidate models included all first-order predictor combinations of *Turbinaria* %N, water column nutrients (nitrite + nitrate, phosphate, and silicate), and fDOM indices (HIX and BIX). Repeated microbial community structure across years was assessed using Mantel tests.

All analyses were performed in R version 4.2.2.

## Results

### Spatial variation in nutrient enrichment

Long-term measures (2007–2022) of *Turbinaria* %N content from three habitats in each of six MCR LTER sites reveal consistent patterns in nutrient loading among habitat types, with elevated *Turbinaria* %N evident near shorelines in fringing reef habitats compared to back reef and reef crest habitats located further offshore (Fig. 1c). This pattern was consistent among each of the six sites (Supplementary materials S7). Across the long-term MCR LTER dataset, *Turbinaria* %N was highest at fringing sites (p < 0.001) but did not significantly differ among back reef and reef crest sites (p = 0.3). Nutrient enrichment was positively associated with land clearing, and 2/3 of the variability of overall mean *Turbinaria* %N was explained by distance from shore and cleared area in a simple linear model (R^2^ = 0.63, p < 0.001). Within the level of individual habitat types, land clearing and distance from shore explained over 90% of the variability of *Turbinaria* %N (p = 0.01) for fringing reef sites, but these factors were not significantly associated with *Turbinaria* %N for back reef or reef crest sites (p = 0.7 and 0.8, respectively).

The finer spatial scale analysis using the 204 lagoon sampling locations highlighted the high spatial heterogeneity of both *Turbinaria* %N and water column N concentration around the island. *Turbinaria* %N significantly varied by habitat type (p < 0.001), with the highest %N at fringing reef sites and in bays and their passes, and lower %N at mid-lagoon and back reef sites (Fig. 2a). Water column N also varied by habitat (p < 0.001), showing a similar pattern of variation, with highest nitrite + nitrate in bays, followed by fringing reef, reef pass, and finally mid lagoon and back reef sites (Fig. 2b). In general, *Turbinaria* %N was positively associated with water column N (Fig. 2c; R^2^ = 0.14, p < 0.001). Mean *Turbinaria* %N was highest near shorelines and declined linearly with increasing distance from shore (Fig. 2d; R^2^ = 0.10, p < 0.001). Residual *Turbinaria* %N variability around the distance-from-shore relationship was positively associated with cleared area of upstream watersheds (p = 0.02), revealing nutrient hotspots at several reef passes and three predominant bays: the bay at Atiha near the southwest tip of the island, Opunohu Bay on the north shore, and the bay at Afareaitu on the east shore (Fig. 2e, Supplementary materials S8).

**Figure 2.**
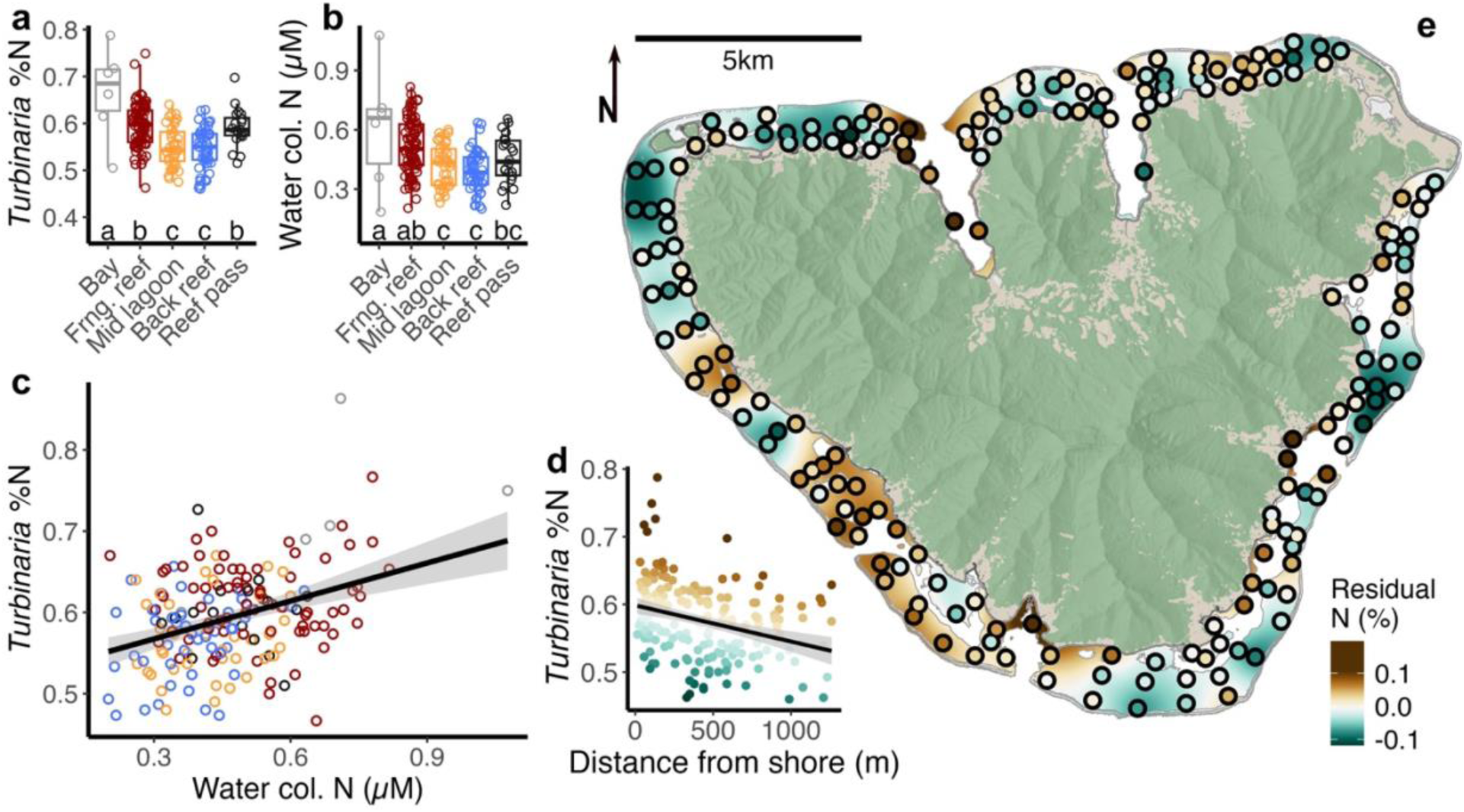
Nutrient indices from *Turbinaria* tissue and water column samples were elevated in bays (grey), fringing reefs (red), and reef passes (black), and lower N at mid lagoon (yellow) and back reef (green) sites (a, b; annotated letters show significant differences [p<0.05] among habitat after accounting for interannual differences using estimated marginal means). Across sites, mean *Turbinaria* tissue %N and water column N measurements were positively correlated (c; p<0.001). The overall mean, gap-filled *Turbinaria* %N dataset showed a negative association with distance from shore (d; p<0.001). Krigged residual values from the distance-from-shore association of mean *Turbinaria* %N (color ramp in d and e) reveal areas with unexpected *Turbinaria* %N given their location (e) with areas in brown showing higher and areas in green showing lower *Turbinaria* %N than expected, respectively.

### Patterns of water chemistry associated with habitat and nutrients

Several fDOM components (ultraviolet humic-like, visible humic-like, and marine humic-like), fDOM indices (HIX, FI, and M:C), and water column nutrients (nitrite + nitrate, phosphate, and silicate) differed by habitat (p < 0.05, Supplementary materials S9). Water column nutrients were highest in bays and fringing reefs, and lowest in mid-lagoon and back reef sites, and reef passes had intermediate silicate and nitrite + nitrate compared to other habitats. This pattern was generally maintained for fDOM components and HIX, and inversely, for BIX, FI, and M:C with varying degrees of significance (Supplementary materials S9). Every water column chemistry parameter (i.e. all water column nutrients, fDOM components, and fDOM indices in this analysis) differed significantly by year (p < 0.05). With the exception of M:C, interactions between habitat and year were never significant for any of the water chemistry parameters.

A principal components analysis showed that water column nutrients (namely nitrite + nitrate and silicate) were often associated with one another, and water column fDOM components (humics and the larger lignin-like, tryptophan-like, and tyrosine-line macromolecules) were often associated with one another (Fig. 3). Quantities of fDOM components were orthogonal to those of nutrients, whereas indices of composition generally covaried with nutrients, suggesting there were additional sources of fDOM in the reef independent of terrestrial inputs that covaried with nutrients. PERMANOVA revealed significant differences in the water chemistry community by discrete habitat type (Fig. 3a, p = 0.001). Further, overall mean *Turbinaria* %N (p = 0.001), distance from shore (p = 0.001), residual *Turbinaria* %N after accounting for distance from shore (p = 0.001), and the area of cleared land in the upstream watershed (p = 0.001) were all significantly associated with the first two water chemistry principal components (Fig. 3b). Notably, *Turbinaria* %N and the amount of cleared land loaded positively in association with water column N, while distance from shore loaded negatively in association with water column N. Within individual years, water column nutrients were consistently positively associated with *Turbinaria* %N and negatively associated with distance from shore every year, and also positively associated with cleared area of the upstream watershed in 2021 (Supplementary materials S10; p < 0.01 in all cases).

**Figure 3.**
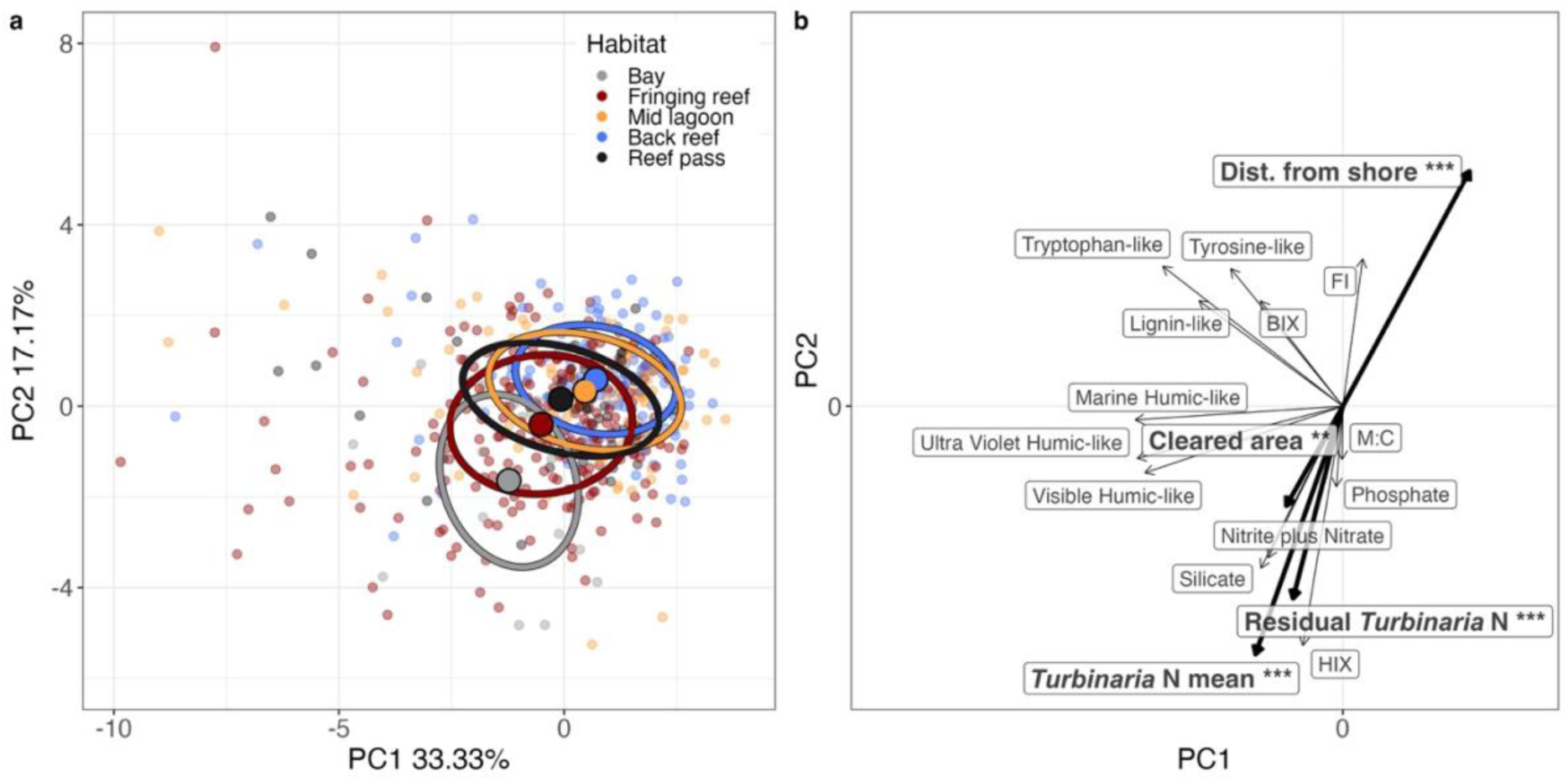
Ordination of water column nutrients and fluorescent dissolved organic matter using principal component analysis. PERMANOVA reveals significant differences in water column chemistry among habitat types (a; R^2^=0.08, *F*=12.45, p=0.001; ellipses based on 1 standard deviation, and centroids shown as bold dots). Generalized additive models identify significant associations between continuous environmental parameters and the ordination space (b; bold arrows; r^2^=0.15, 0.03, 0.15, and 0.09 for Distance from shore, Cleared Area, *Turbinaria* %N, and detrended *Turbinaria* %N after accounting for distance from shore respectively, and p<0.01 in all cases).

### Effects of recent precipitation on lagoon nutrients

Cumulative precipitation had different effects on lagoon nutrients over short and long timescales. Water column N was positively associated with cumulative precipitation over short timescales (<5 days) but negatively associated with cumulative precipitation over longer timescales (Fig. 4a). Conversely, *Turbinaria* %N was negatively associated with cumulative precipitation over short timescales but positively associated with cumulative precipitation over longer timescales (Fig. 4a). Linear mixed-effects models revealed that lagoon nutrients (indexed both by *Turbinaria* %N and water column N) were consistently positively associated with cleared land in upstream watersheds regardless of the timescale of cumulative precipitation considered in the analysis (Fig. 4b).

**Figure 4.**
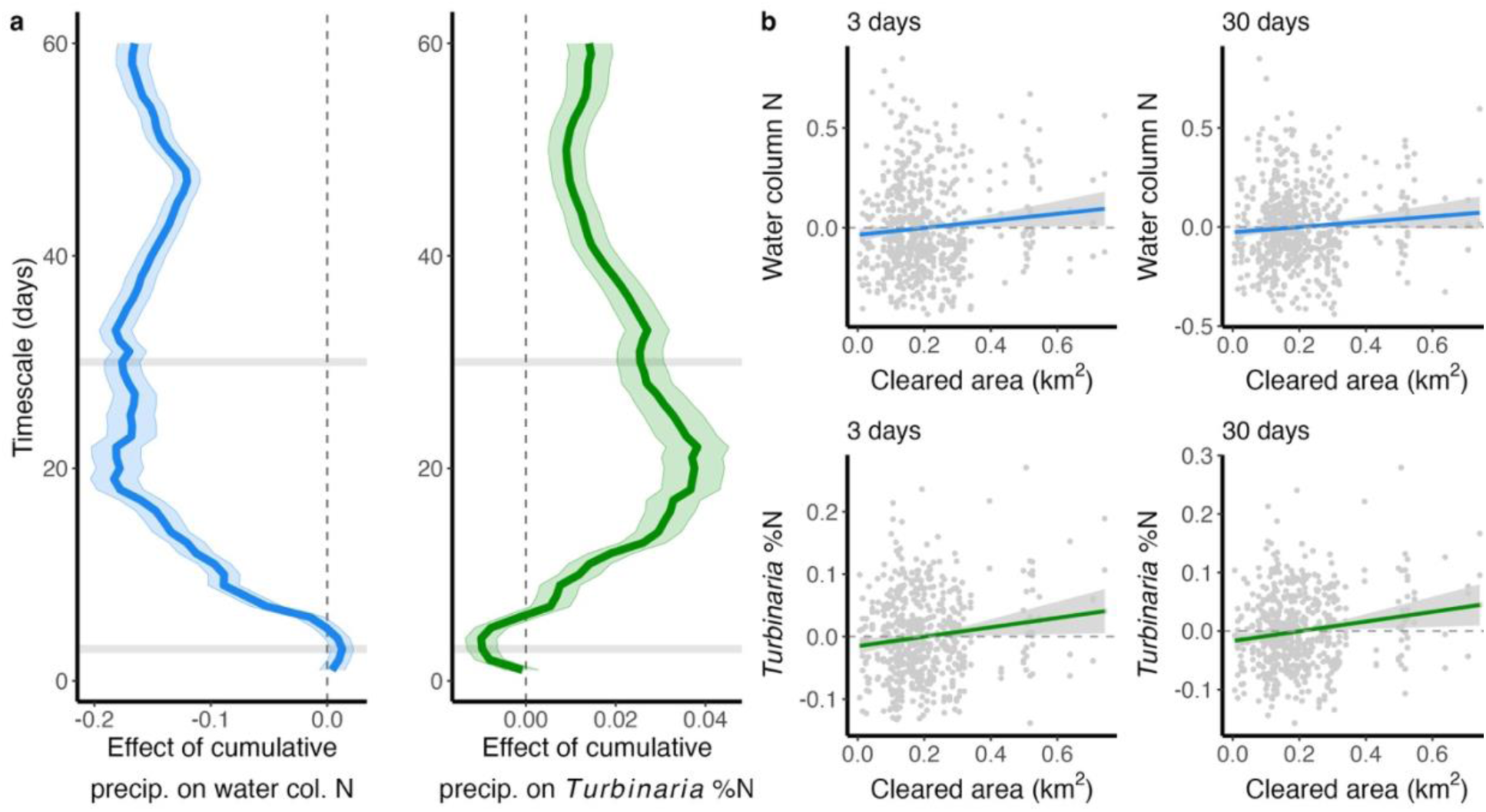
The role of recent cumulative precipitation in lagoon nutrient dynamics depends on timescale (a). Over short timescales, water column N (blue) increases with increasing recent rain, but over longer timescales, water column N is negatively associated with cumulative precipitation. Conversely, *Turbinaria* %N (green) is negatively associated with cumulative precipitation over short timescales, but positively associated with cumulative precipitation over longer timescales. Cleared area of upstream watersheds is associated with increased N in both water column and *Turbinaria* samples (b; shown are contrast plots with partial residuals), after controlling for recent cumulative precipitation regardless of timescale (left, 3 days; and right, 30 days).

### Associations between microbial communities and nutrient regimes

Microbial communities differed by habitat (Fig. 5a, PERMANOVA, p = 0.001). While mid lagoon, back reef, and reef pass sites had similar microbial communities (indicated by highly overlapping regions in PCoA space), fringing reefs and bays featured divergent microbial communities. An analysis of homogeneity of group dispersions showed that fringing reefs had more variable communities than mid-lagoon and back reef sites, and bays and reef passes had more variable communities than backreef sites (p < 0.05 in all cases). Together, distance from shore and residual *Turbinaria* %N explained over 1/3 of the variability in microbial communities along MDS 1 (p < 0.01). Notably, sites that were close to shore had microbial communities more similar to offshore sites when nutrients were low, and more disparate microbial communities when nutrients were high (Fig. 5b). Based on the model selection framework, patterns in microbial communities were best explained by *Turbinaria* %N and water column nitrite + nitrate, silicate, and HIX (Table 1; p < 0.01 in all cases). Mantel tests on Bray-Curtis distance matrices revealed consistent spatial structure of lagoon microbial communities among years (p = 0.001 in all 3 comparisons; Mantel r = 0.45, 0.52, and 0.44 for 2021 vs 2022, 2021 vs 2023, and 2022 vs 2023, respectively; Supplementary materials S11).

**Figure 5.**
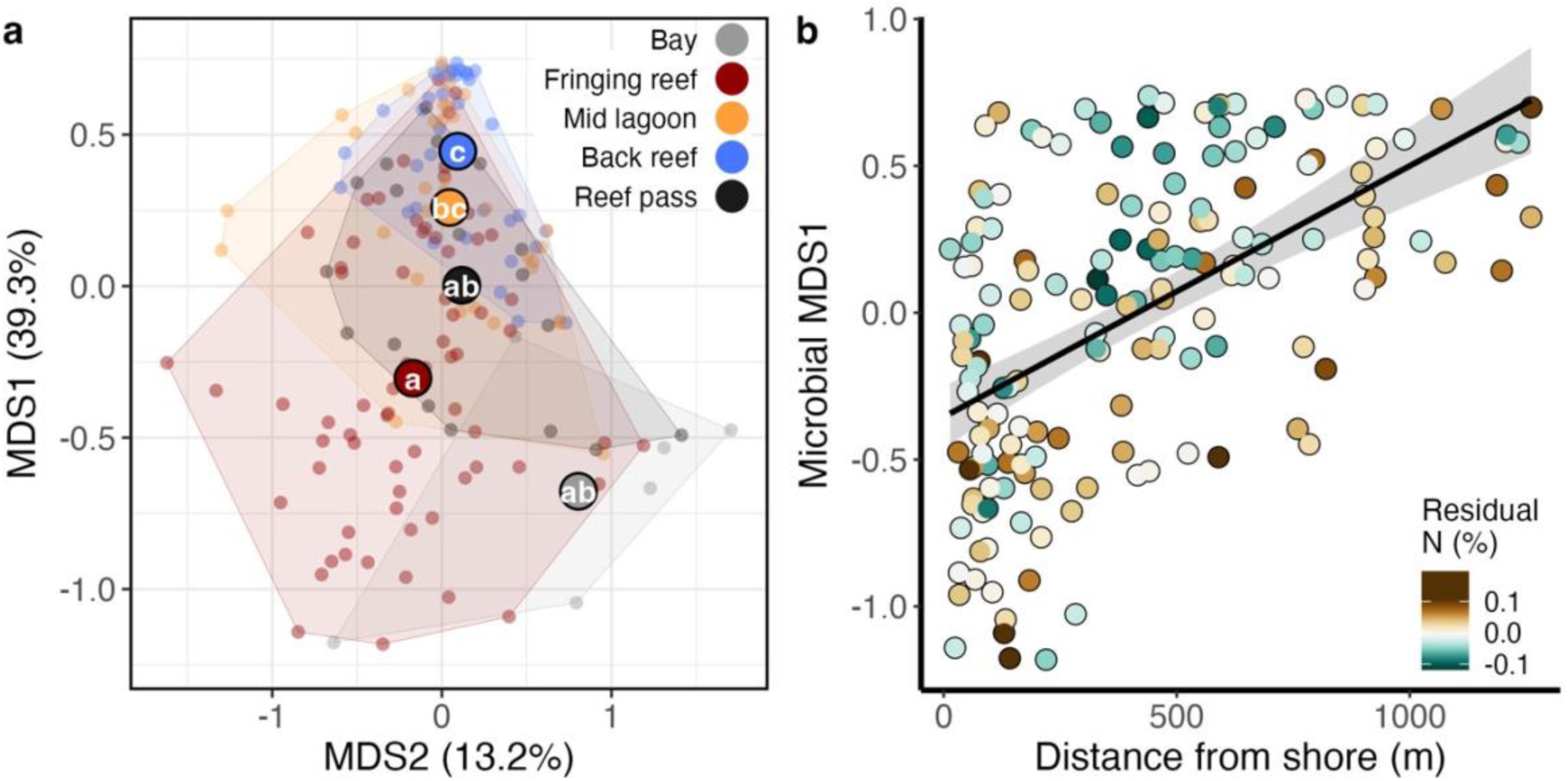
Ordination of water column microbial communities using PCoA with Bray-Curtis distance, aggregated to the genus level (a). Points are colored by habitat type (habitat centroids shown as bold points). Comparisons of microbial community dispersion by habitat showed that back reef sites were less dispersed than bays, fringing reefs, and reef pass sites, and mid-lagoon sites were less dispersed than fringing reef sites (letters annotated in a). Variation in microbial community composition along MDS axis 1 was explained by distance from shore and residual variation in *Turbinaria* %N after accounting for distance from shore (b).

**Table 1.** Summary of PERMANOVA results for selected environmental variables explaining dissimilarity among microbial communities.

| *term* | *Df* | *Sum of squares* | *R*<sup>2</sup> | *F* | *p* |  |
| --- | --- | --- | --- | --- | --- | --- |
| *Turbinaria* %N | 1 | 1.01 | 0.09 | 18.96 | 0.001 | *** |
| Nitrate plus Nitrite | 1 | 0.55 | 0.05 | 10.39 | 0.001 | *** |
| Silicate | 1 | 0.71 | 0.06 | 13.34 | 0.001 | *** |
| HIX | 1 | 0.33 | 0.03 | 6.22 | 0.001 | *** |
| Residual | 160 | 8.51 | 0.77 |  |  |  |
| Total | 164 | 11.12 | 1 |  |  |  |

## Discussion

Nutrient availability is an important driver of community structure in coastal ecosystems in the tropics as they are often oligotrophic. Nutrient subsidies could modify spatial patterns of community dynamics if nutrient sources are spatially structured. In this study, we combined a fine-scale sampling array of tissues from an algal bioindicator of nitrogen enrichment, with water column chemistry, land use, and precipitation to uncover drivers of lagoon nutrient regimes and their influence on microbial community dynamics in a tropical lagoon ecosystem. We showed that long-term nutrient dynamics reflect fine-scale variability in water column chemistry across shorter temporal scales. Water column nutrients and terrigenous organic matter decreased with increasing distance from the island and increased when nearby watersheds had more cleared land area. Whereas water column N was elevated following rainfall on 3-day timescales, tissue N content of *Turbinaria* was elevated following rainfall over monthlong timescales. Finally, unique microbial communities were found in the habitats most proximal to terrestrial impacts: fringing reefs and bays.

### Patterns of lagoon nutrient regimes

Tissue N content of the brown macroalga *Turbinaria ornata* (*Turbinaria* %N) was highest in bays, fringing reefs and reef passes, and lowest at mid lagoon and back reef sites. Back reef sites, which are ostensibly most isolated from terrestrial impacts, consistently featured the lowest *Turbinaria* %N across shores. We identified a similar pattern of variation in water column samples, with the lowest N concentrations at mid lagoon and back reef sites. Nutrients can be introduced to nearshore systems from several sources. Land-based nutrient sources include runoff, wastewater, and submarine groundwater discharge (SGD) (Dollar and Atkinson 1992; Hernández-Terrones et al. 2015; Barnas et al. 2025; Silbiger et al. 2025) while oceanic inputs include offshore currents, swell, upwelling, and wave action (Pfister et al. 2007; Woodson 2018). Our finding that nutrients were consistently highest close to shore indicate that nutrients in Moorea lagoons are enriched by land-based sources, declining with increasing distance from shore due to biological uptake of nutrients or dilution mechanisms (Lapointe and Clark 1992; Abaya et al. 2018). Residual values from the relationship between *Turbinaria* %N and distance from shore reveal that nutrient enrichment hotspots around Moorea often occurred at reef passes, which had higher N than mid lagoon or back reef sites (residual %N at reef pass sites was higher by 0.031% and 0.024% than mid-lagoon and back reef sites, respectively). We attribute this to increased flux of nutrients from nearby bays, which are especially prone to accumulations of terrigenous matter (Dame et al. 1986; Howarth et al. 2000), due to high water movement through the passes resulting in increased N accumulated in algal tissues despite their farther distances from shore.

Importantly, we identified a weak but positive association between *Turbinaria* tissue %N and water column N. Direct measurements of water column chemistry and indirect measurements via bioindicators provide complementary perspectives of nutrient availability; whereas water column samples provide instantaneous measures of lagoon nutrients, algal bioindicators lend insights on nutrient dynamics over weeks-to-months’ timescales (Salo and Salovius-Laurén 2022). Nutrient contents of *Turbinaria* tissues have been used as a proxy for nutrient enrichment over weeks to months-long timescales in this system and elsewhere (Schaffelke 1999; Donovan et al. 2020; Adam et al. 2021), but short-term environmental variability such as precipitation or ephemeral water currents are important drivers of lagoon water chemistry dynamics in Moorea (e.g. Fong et al. 2020). Whereas water column attributes may be fleeting, longer time-integrated measurements from *Turbinaria* tissues reflect accumulated differences that may not be evident over short timescales. Our results indicate that despite short-term environmental variability, nutrient regimes summarized by a bioindicator can be a meaningful proxy for chronic nutrient enrichment.

Mean %N content of *Turbinaria* tissues was positively associated with multiple water column nutrient and fDOM parameters in our water chemistry ordination, further highlighting the utility of *Turbinaria* %N as an indicator for water quality. Specifically, *Turbinaria* %N was positively associated with water column nitrite + nitrate, silicate, HIX, and, negatively associated with FI. Silicate is associated with submarine groundwater discharge, suggesting that SGD is a potential pathway of nutrient enrichment in the lagoons of Moorea (Nelson et al. 2015; Oehler et al. 2019). Lower FI values indicate that water column organic matter derives from terrestrial origins (Cory and McKnight 2005), and higher values of HIX indicate a higher total load of humics in the sample (Zsolnay et al. 1999). Together, the positive correlation between water column N and HIX, the negative correlation between water column N and FI, and the positive correlation between FI and distance from shore, corroborate our conclusion that there are significant terrigenous sources of nutrients to the lagoons of Moorea.

### Dynamic influence of terrestrial factors on lagoon nutrients

Our results showed that precipitation played a dynamic role in influencing patterns of lagoon nutrients, and that the relationship between lagoon nutrients and participation varied by timescale. Whereas water column N was positively correlated with cumulative precipitation over short timescales (<5 days), it was negatively correlated with cumulative precipitation over longer timescales. In contrast, *Turbinaria* %N showed the opposite pattern. We suspect that the comparative drought conditions during the month leading up to sampling in 2022, followed by extreme rainfall in the last few days prior to that sampling event, led to an accretion of soluble nutrients in soils which was suddenly released at the point of the most recent precipitation event (Szejgis et al. 2024). In the other years, consistent rain during the months leading up to sampling could have driven regular runoff of surface nutrients which had depleted by the time water column sampling occurred. Thus, positive associations between cumulative rain and *Turbinaria* %N over long timescales indicates that chronic N enrichment delivered by precipitation runoff is reflected in the bioindicator nutrient record but missed by instantaneous water column measurements.

Furthermore, we found that nutrient enrichment was positively associated with land clearing in nearby watersheds. Regardless of the timescale of precipitation impacts, *Turbinaria* %N and water column nitrite + nitrate were higher downstream of watersheds with increased land clearing. Alongside the positive association between these factors and HIX, these results point toward land use as an important source of variation in terrestrial subsidies for lagoon nutrients. Previous work has shown that land clearing on Moorea leads to increased sediment and nutrient content in streams, and this association is related to recent precipitation (Neumann et al. 2025). Our results point toward a broader downstream impact of land use on aquatic nutrient dynamics, particularly in habitats that are close to shore. Further work should explore how riparian corridors and phytoremediation could temper this effect (Carlson et al. 2019), and identify how nitrogen-fixing forest plants like the prevalent and invasive *Falcataria moluccana* drive additional variability in nutrient subsidies from land (Atwood et al. 2010).

### Lagoon nutrients shape planktonic microbial communities

We found that nutrient regimes (summarized by mean *Turbinaria* %N) were significantly correlated with microbial community structure. Whereas we expected microbial communities in different reef habitats to be unique (Leichter et al. 2013; Hamamoto et al. 2024), communities in habitats more isolated from terrestrial influence – i.e. mid lagoon and back reef sites – had similar microbial communities and even overlapped with a subset of fringing reef sites. However, in habitats closest to the island – i.e. bays and fringing reef sites – microbial communities diverged relative to other lagoon habitats. Communities in these bay and fringing reef sites departed from those of other lagoon habitats in association with increasing *Turbinaria* %N. Microbial community diversity along nearshore-offshore gradients has been linked to abiotic factors including temperature and nutrients over broad scales (10’s – 100’s of km) (Wang et al. 2019; James et al. 2022). The unique microbial communities found in N-enriched bay and fringing reef habitats of Moorea suggest that abiotic variability is also an important factor driving onshore-offshore patterns of microbial community structure at fine scales (100’s of m) (Nelson et al. 2011; Leichter et al. 2013).

### Conclusions and future directions

Changing nutrient regimes in other nearshore systems have been attributed to interactions among climatic factors and modified terrestrial inputs under changing human activities (Xin et al. 2019; Brodie et al. 2020). Ensemble models suggest that the South Pacific will be warmer and wetter in the coming decades (Ying et al. 2022). Therefore, Moorea’s future lagoon nutrient dynamics will hinge upon links between land use and precipitation, and the implementation of catchment management strategies that account for fine-scale complexity. Here, we explored how terrestrial factors influence nearshore biogeochemical and microbial community dynamics across the lagoons of Moorea. We found that nutrient enrichment was associated with land use and mediated by precipitation and identified nutrients as a potential link between these factors and differences in planktonic microbial communities. Future work should investigate direct associations between humans, nutrients, and microbes, to better understand the pathways by which nutrients are introduced to nearshore systems and causal associations between water chemistry and microbial communities. Developing mechanistic connections between human activities, climate change, and lagoon nutrients will help inform management decisions that minimize terrestrial impacts on sensitive coral reef lagoon systems.

## Acknowledgements

The authors are grateful to the staff of Gump Research Station for logistical support. This research was supported by the Zegar Family Foundation, Oceankind Inc., and the U.S. National Science Foundation’s (NSF) Moorea Coral Reef Long Term Ecological Research (MCR LTER) site under Grant OCE 2224354 (and earlier awards). Additional financial support to the MCR LTER site was provided through a generous gift from the Gordon and Betty Moore Foundation. Research was also supported by the Uehiro Center for Advancement of Oceanography and NSF-OCE #1924281 to Nyssa J Silbiger. This is contribution # XXX (SOEST), XXX (UC•AO), and XXX(CSUN). Research was completed under permits issued by the French Polynesian Government (Délégation à la Recherche) and the Haut-commissariat de la République en Polynésie Francaise (DTRT) (Protocole d’Accueil 2005–2025). With respect to the spelling of Moorea, we followed the Raapoto transcription system that is adhered to by a large segment of the Tahitian community, but also recognize other community members follow the Te Fare Vāna’a transcription system where the island name is spelled with an ’eta (Mo’orea).

## Author contributions

Conceptualization: DEB, MKD, CJ, CEN, NJS, RVT

Data curation: TCA, DEB, RC, MKD, CJ, CEN, DPS, RVT

Formal analysis: CJ, NJS, CEN, MKD

Funding acquisition: DEB, NJS, RVT, TCA, CEN

Investigation: all authors

Methodology: TCA, CJ, CEN, NJS, DEB, RVT

Project administration: DEB, TCA, RVT

Software: CJ, NJS

Visualization: CJ

Writing – original draft: CJ, NJS, TCA, MKD, LWK, CEN, RVT, DEB

Writing – review and editing: all authors

## Data availability statement

And all data are available on the Environmental Data Initiative repository (links and DOI’s will be provided upon acceptance for publication***) and code to reproduce results is available on github (links will be provided upon acceptance for publication***).

