## Supplementary materials for "Terrigenous inputs link nutrient dynamics to microbial communities in a tropical lagoon"

Supporting information for:
Terrigenous inputs link nutrient dynamics to microbial communities in a tropical lagoon

**Terrigenous inputs link nutrient dynamics to microbial communities in a tropical lagoon**

**Authors:**
Christian John[1*], Nyssa J. Silbiger[2], Thomas C. Adam[3], Danielle M. Barnas[4], Kalia S. Bistolas[5], Robert C. Carpenter[6], Noe Castañeda[7], Megan J. Donahue[8], Mary K. Donovan[9], Lauren N. Enright[10], Hannah E. Epstein[11], Jordan P. Gallagher[12], Hendrikje Jorissen[13], Jamie R. Kerlin[14], Savanah L. Leidholt[15], Rowan H. McLachlan[16], Nury Molina[17], Catherine A. Mullenmeister[18], Kyle Neumann[19], Julianna J. Renzi[20], Denise P. Silva[21], Kelly E. Speare[22], Sean Swift[23], Alex D. Vompe[24], Linda Wegley Kelly[25], Maya Zeff[26], Craig E. Nelson[27], Rebecca Vega Thurber28, Deron E. Burkepile[29]

**Affiliations & ORCID ID's:**
[1]Marine Science Institute; University of California, Santa Barbara.
[2]Uehiro Center for the Advancement of Oceanography, Department of Oceanography; University of Hawaiʻi at Mānoa.
[3]Marine Science Institute; University of California, Santa Barbara.
[4]Hawaiʻi Institute of Marine Biology; University of Hawaiʻi at Mānoa.
[5]Department of Microbiology; Oregon State University.
[6]Department of Biology; California State University, Northridge.
[7]Department of Ecology, Evolution and Marine Biology; University of California, Santa Barbara.
[8]Hawaiʻi Institute of Marine Biology; University of Hawaiʻi at Mānoa.
[9]Hawaiʻi Institute of Marine Biology; University of Hawaiʻi at Mānoa.
[10]Department of Ecology, Evolution and Marine Biology; University of California, Santa Barbara.
[11]School of Life Sciences; University of Essex. Department of Microbiology; Oregon State University.
[12]Department of Ecology, Evolution and Marine Biology; University of California, Santa Barbara.
[13]Hawaiʻi Institute of Marine Biology; University of Hawaiʻi at Mānoa.
[14]Department of Biology; California State University, Northridge.
[15]Department of Microbiology; Oregon State University.
[16]Oregon Institute of Marine Biology; University of Oregon.
[17]Department of Ecology, Evolution and Marine Biology; University of California, Santa Barbara.
[18]Marine Biology Research Division, Scripps Institution of Oceanography; UC San Diego.
[19]Marine Science Institute; University of California, Santa Barbara.
[20]Department of Ecology, Evolution and Marine Biology; University of California, Santa Barbara.
[21]Marine Science Institute; University of California, Santa Barbara. Department of Microbiology; Oregon State University.
[22]Marine Science Institute; University of California, Santa Barbara. School of Geographical Sciences and Urban Planning; Arizona State University.
[23]Daniel K. Inouye Center for Microbial Oceanography: Research and Education; University of Hawaiʻi at Mānoa. Sea Grant College Program; University of Hawaiʻi at Mānoa.
[24]Department of Plant Pathology and Environmental Microbiology; The Pennsylvania State University.
[25]Marine Biology Research Division, Scripps Institution of Oceanography; UC San Diego.
[26]Department of Ecology and Evolutionary Biology; University of California, Santa Cruz.

Supporting information for:
Terrigenous inputs link nutrient dynamics to microbial communities in a tropical lagoon

[27]Daniel K. Inouye Center for Microbial Oceanography: Research and Education; University of Hawaiʻi at Mānoa. Department of Oceanography and Sea Grant College Program; University of Hawaiʻi at Mānoa.
[28]Marine Science Institute; University of California, Santa Barbara.
[29]Marine Science Institute; University of California, Santa Barbara. Department of Ecology, Evolution, & Marine Biology; University of California, Santa Barbara.

***Corresponding author:**
Christian John

Marine Science Institute
University of California, Santa Barbara
Santa Barbara, CA  93106, USA

Supporting information for:
Terrigenous inputs link nutrient dynamics to microbial communities in a tropical lagoon

**Supplementary materials S1: Sampling metrics from island-wide sampling campaign**
Sampling effort (number of sites) for *Turbinaria* tissues, water column nutrients, water column fDOM, and planktonic microbial communities from fine-scale island-wide sampling campaign.

| Dataset | May 2016 | May 2021 | April 2022 | April 2023 |
|---|---|---|---|---|
| *Turbinaria* %N | 171 | 193 | 190 | 198 |
| Water nutrients | - | 195 | 191 | 200 |
| Water fDOM | - | 194 | 173 | 195 |
| Microbes | - | 195 | 187 | 167 |

**Supplementary materials S2: Intercomparison of bioindicator at different analytical labs**
Cross-correlation of laboratory analyses of *Turbinaria* tissue %N from 2016 samples sent to both NA1500 (Carlo-Erba) at UGA and CEC440HA (Exeter Analytical) at UCSB (n = 20).

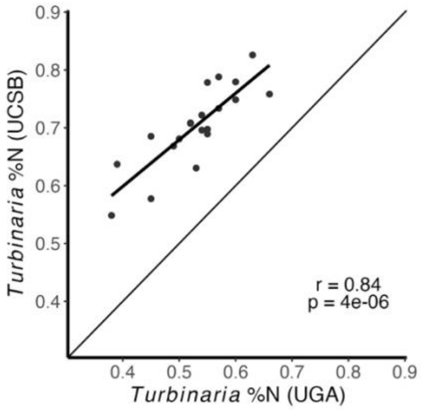

Supporting information for:
Terrigenous inputs link nutrient dynamics to microbial communities in a tropical lagoon

**Supplementary materials S3. Additional details of water chemistry sample processing**

Water column nutrients were analyzed using flow injection with an automated ion analyzer (QuickChem 8500, Zellweger Analytics). In this instrument, samples are passed through narrow-gauge tubing manifolds and combined with reactants under controlled diffusion rates and can be analyzed for multiple nutrients and other ions simultaneously. Analyte concentrations were determined using the Sulfanilamide/NED Cd reduction method 31-107-04-1-A at 520nm for nitrite + nitrate (detection limit = 0.2µM), the molybdate-based method 31-115-01-1-A at 880nm for Phosphate (detection limit = 0.1µM), and the Molybdate reactive method 31-114-27-1-B at 820nm for Silicate (detection limit = 1.0µM) (Lachat Instruments 2836; precision = ±5% for all analytes).

For water column fDOM, water samples were stored at 4 °C and brought to room temperature one hour prior to analysis. 3 mL subsamples were transferred into 1 cm quartz cuvettes, which were then loaded into a Horiba Aqualog scanning fluorometer. Prior to running samples, cuvettes were washed thoroughly with deionized water (DIW) and wiped clear with Kim wipes. Cuvettes were triple rinsed with DIW between each sample. After every 7 samples, a DIW blank was run. Excitation-Emission Matrices (EEMS) were generated with excitation values ranging from 240 nm to 500 nm, and emission values were measured ranging from 250 nm to 825 nm. Excitation wavelengths were increased at 5 nm increments, and measurements were integrated over 4 s at each increment. Inner filter correction was applied to account for fluorescence quenching by absorbance (Kothawala et al. 2013). Fluorescence values were scaled to Raman Units by dividing by the integrated emission range of 381 nm to 426 nm at an excitation of 350 nm in averaged DIW blanks (Lawaetz and Stedmon 2009). The average blank EEMs, from DIW blanks run immediately before and after each set of 7 samples, were subtracted from the sample EEMs.

Supporting information for:
Terrigenous inputs link nutrient dynamics to microbial communities in a tropical lagoon

**Supplementary materials S4. Additional details of water column microbial community sample processing**

Modified DNA extraction as follows: Sterivex filter units were defrosted and sealed at one end using a Luer lock cap (red cap TrueCare, South Miami, Florida, USA), then 440µL of extraction buffer solution (40µL proteinase K, 200µL of buffer AL provided by kit, and 220µL of PBS; [Gibco, New York, USA]) was added to the column and sealed with another Luer cap, and then each end was wrapped in parafilm to avoid leakage. Filled filter units were attached horizontally to the rotator in a hybridization incubator and incubated at 56°C for 4 hours while rotating at 20 rpm. After incubation, the inlet cap was removed, and the inlet port of the filter unit was added to the 2 mL tube and sealed with parafilm. The filter cartridge with a 2mL tube was allocated into the sterile 50mL conical tube and centrifuged at 5,000 x g for 2 min to collect the extracted DNA from the filter cartridge. Next, the 2mL tube was separated from the filter cartridge, and 200µL 100% ethanol was added to the extracted DNA and mixed thoroughly by vortexing. The mixture was then transferred to a DNeasy kit and processed according to the manufacture recommendations.

One-step PCR consisted of 12.5µl AccuStart II PCR ToughMix (2x) (Quanta BioSciences, Gaithersburg, Maryland, USA), 7.5µl ultra-pure water, 2.5µL forward primer (10 µM), 2.5µl reverse primer (10µM) and 1µl template DNA, totaling 25µL. The thermocycling protocol included an initial denaturation at 94°C for 3 min, followed by denaturation at 94°C for 45s, annealing at 50°C for 60s, and extension at 72°C for 90s. A final extension was performed at 72°C for 10 min, with a final hold at 4°C. The amplified products were visualized by 1.5% agarose gel electrophoresis at 85 volts for 120 minutes. The libraries were purified through the AMPure XP Magnetic Beads (Beckman Coulter Life Sciences, Indiana, EUA). DNA concentrations were quantified using Quant-iT 1X dsDNA High Sensitivity (HS) Assay kit (Thermo Fisher Scientific, Massachusetts, USA).

Supporting information for:
Terrigenous inputs link nutrient dynamics to microbial communities in a tropical lagoon

**Supplementary materials S5. Evaluating effect of missing value interpolation**
Turbinaria tissue samples were collected from 204 unique sites over 4 years. Of the 816 possible site-year sampling events, there were 66 total missing-value cases (i.e. a site could not be visited due to weather conditions, the sample was lost or damaged during processing, etc.). Only sites with at least 3 *Turbinaria* tissue %N measurements (n = 191) were considered; the remaining sites (n = 13) were removed from the analysis. Missing values for retained sites were imputed using a linear model as described in the main text. Of the retained sites, 35 had a missing value that was then imputed. Thus, from the (191 sites * 4 years) 764 total instantaneous *Turbinaria* %N data points, 4.6% were imputed values and 95.4% were observed values. Predicted vs. observed instantaneous *Turbinaria* %N shows strong positive correlation (r = 0.8 and $p < 0.001$).

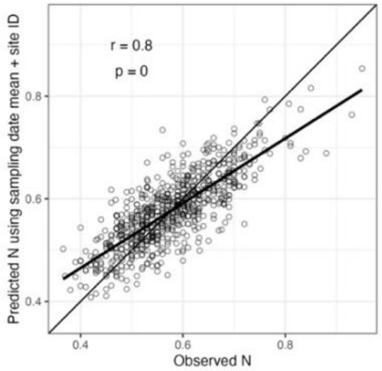


Comparison of main results (Figure 2 main text) with- and without sites containing imputed missing values reveals qualitatively consistent pattern of nutrient enrichment around Moorea. Variability among habitats shows N enrichment at bay and fringing reef sites compared to mid lagoon and back reefs, with reef pass sites at an intermediate level of N enrichment (a and b). *Turbinaria* %N and water column [N] are positively associated across the time series (c). As in the main results, *Turbinaria* %N decreases with increasing distance from shore (d). Removal of sites with imputed values produces a similar krigged N enrichment heatmap (removed sites indicated by grey dots in map, e).

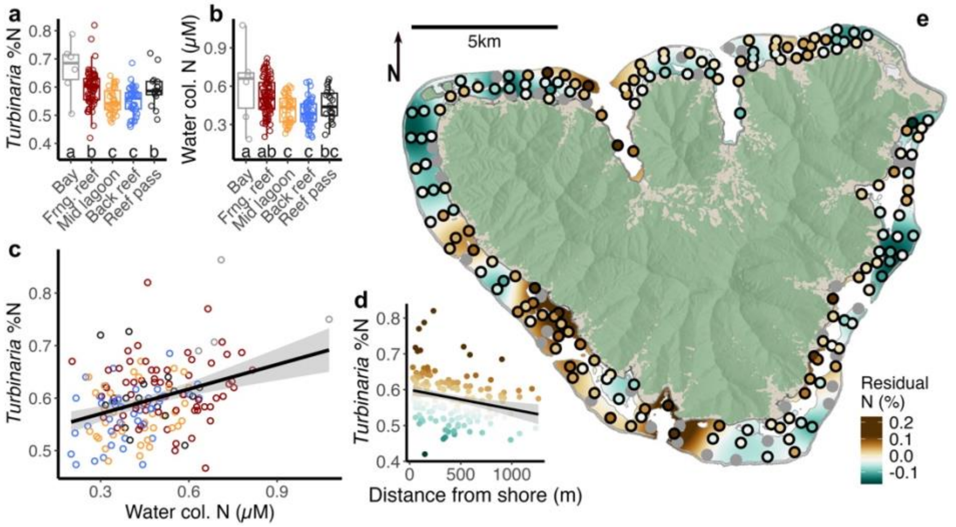

Supporting information for:
Terrigenous inputs link nutrient dynamics to microbial communities in a tropical lagoon

**Supplementary materials S6. Checking assumptions of spherical kriging**
Variogram for the overall spherical kriging approach shows clear plateau and consistent fit around modeled semivariance, indicating stationarity (sum of squared errors = $3.3_{x10}^{-12}$).

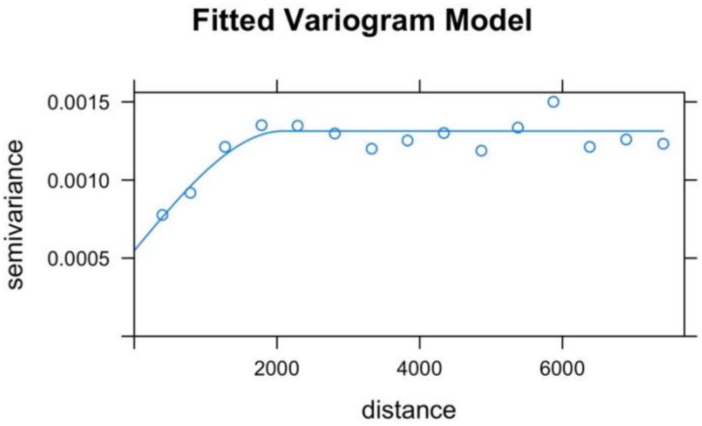


Consistent variogram structure among different point separation directions, indicating isotropy.

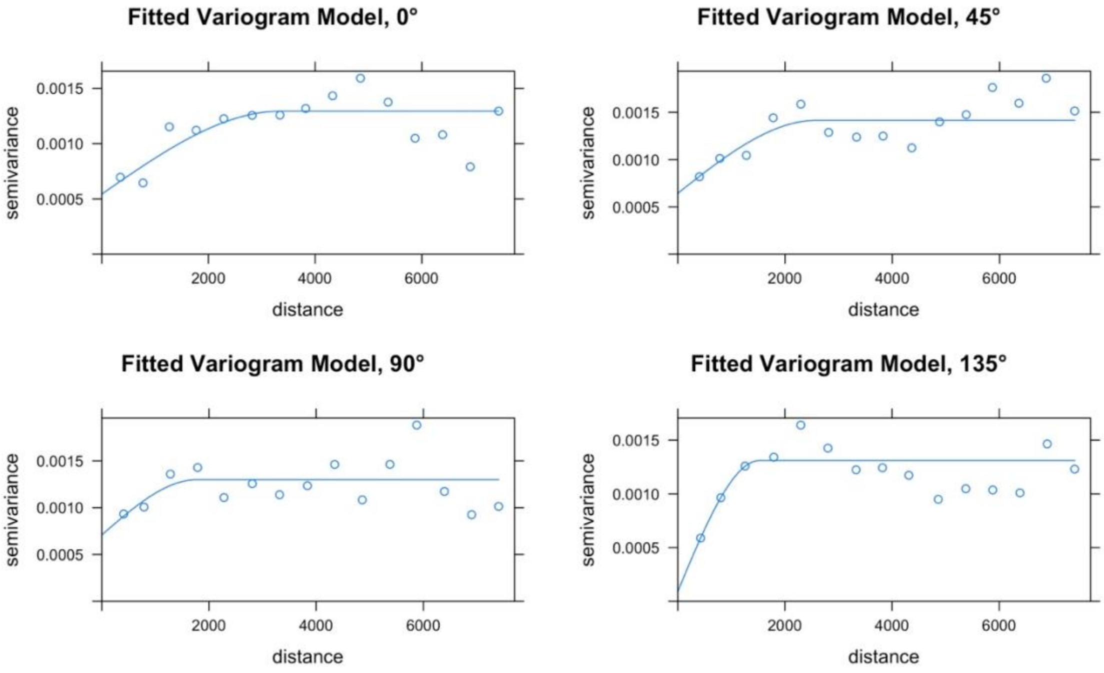

Supporting information for:
Terrigenous inputs link nutrient dynamics to microbial communities in a tropical lagoon

**Supplementary materials S7.** Variability in *Turbinaria* %N measurements from within individual LTER sites shows consistent pattern of variation with overall habitat model presented in main text Figure 1c.

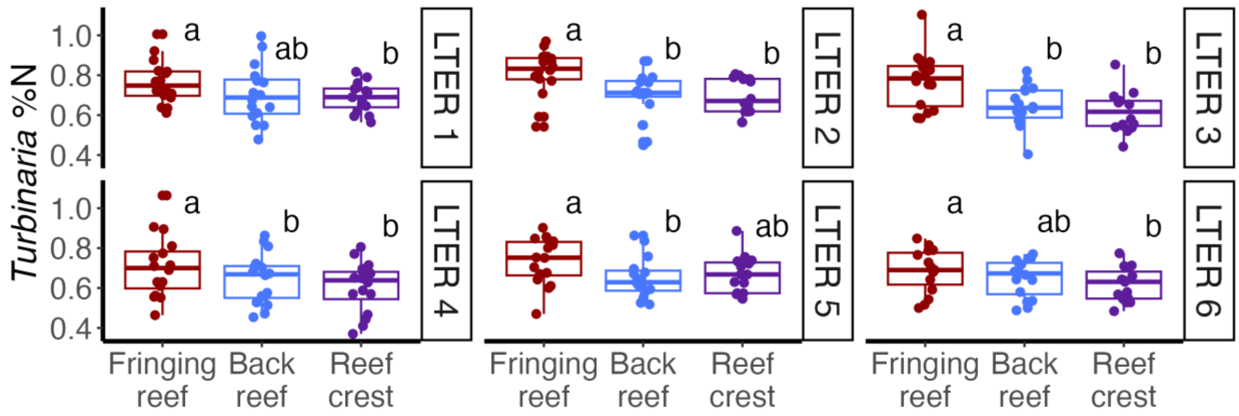

Supporting information for:
Terrigenous inputs link nutrient dynamics to microbial communities in a tropical lagoon

**Supplementary materials S8**. **Residual N variability after accounting for distance from shore**
Residual *Turbinaria* %N was positively associated with cleared area ($p = 0.01$, $R^2 = 0.03$) and not associated with distance from shore.

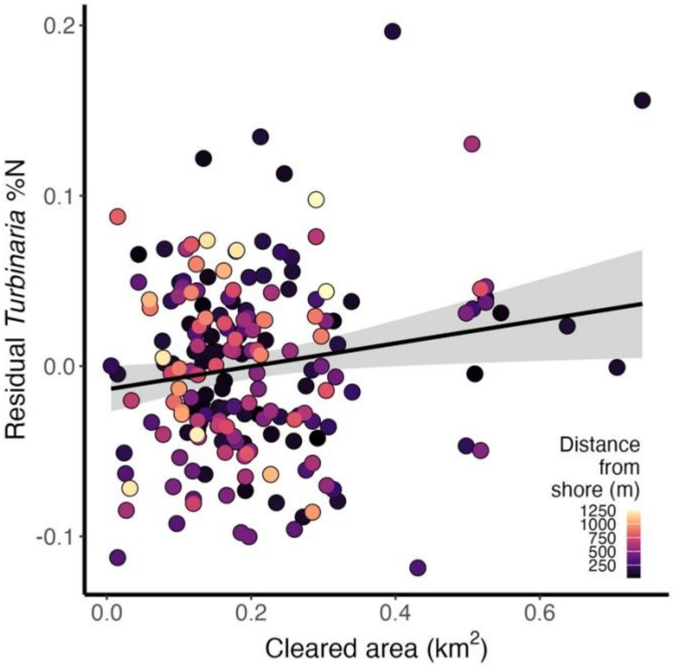


Comparison of residual long-term mean *Turbinaria* %N by habitat shows that Bays and Reef passes had the highest residual N, indicating that in spite of their comparatively large distance from shore, they featured high Turbinaria %N.

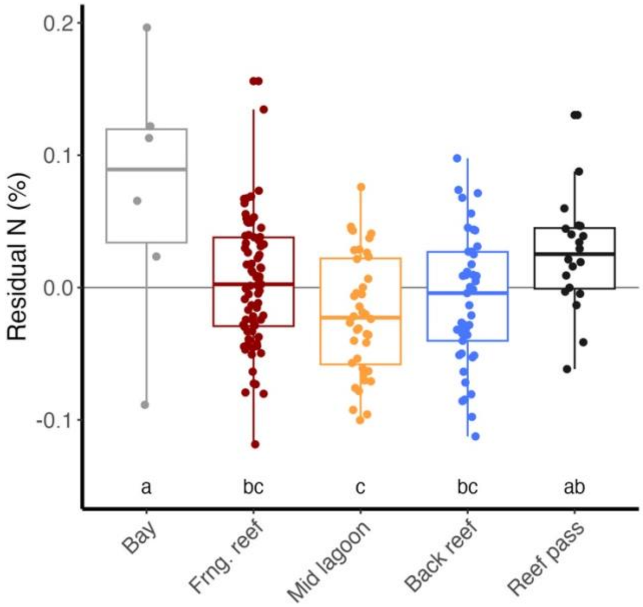

Supporting information for:
Terrigenous inputs link nutrient dynamics to microbial communities in a tropical lagoon

**Supplementary materials S9. Spatiotemporal variability in water column chemistry**
Annual measures by habitat (boxplots) with significant differences among habitats based on estimated marginal means (emmeans) shown as unique letter groups. Emmeans were calculated for linear mixed-effects models with the water column chemistry parameter as the response and habitat as the predictor, and year as a random effect. For tables, summaries are shown for analyses of variance of simple linear models with each water column chemistry parameter as the response and habitat, year, and their interaction as predictors.

**Water column nutrients**

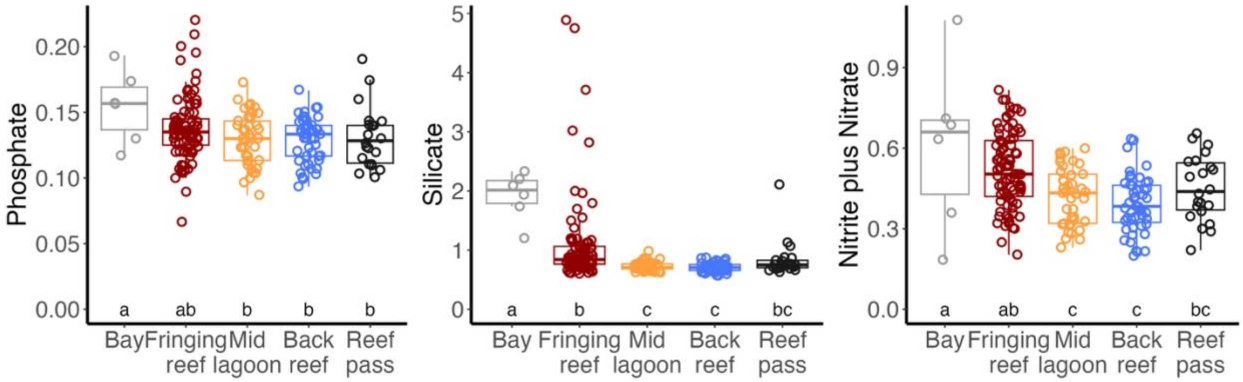


| *Parameter* | *term* | *Df* | *Sum.Sq* | *Mean.Sq* | *F* | *p* | |
|---|---|---|---|---|---|---|---|
| *Nitrate* | Habitat | 4 | 2.37215 | 0.59304 | 11.392 | 0 | *** |
| | Year | 1 | 2.91221 | 2.91221 | 55.94 | 0 | *** |
| | Habitat:Year | 4 | 0.07519 | 0.0188 | 0.361 | 0.836 | |
| | Residuals | 574 | 29.88204 | 0.05206 | | | |
| *Silicate* | Habitat | 4 | 34.91011 | 8.72753 | 16.831 | 0 | *** |
| | Year | 1 | 2.31119 | 2.31119 | 4.457 | 0.035 | * |
| | Habitat:Year | 4 | 0.33049 | 0.08262 | 0.159 | 0.959 | |
| | Residuals | 576 | 298.68409 | 0.51855 | | | |
| *Phosphate* | Habitat | 4 | 0.01698 | 0.00425 | 4.265 | 0.002 | ** |
| | Year | 1 | 0.17171 | 0.17171 | 172.485 | 0 | *** |
| | Habitat:Year | 4 | 0.00673 | 0.00168 | 1.691 | 0.15 | |
| | Residuals | 575 | 0.57243 | 0.001 | | | |

Supporting information for:
Terrigenous inputs link nutrient dynamics to microbial communities in a tropical lagoon

**fDOM components**

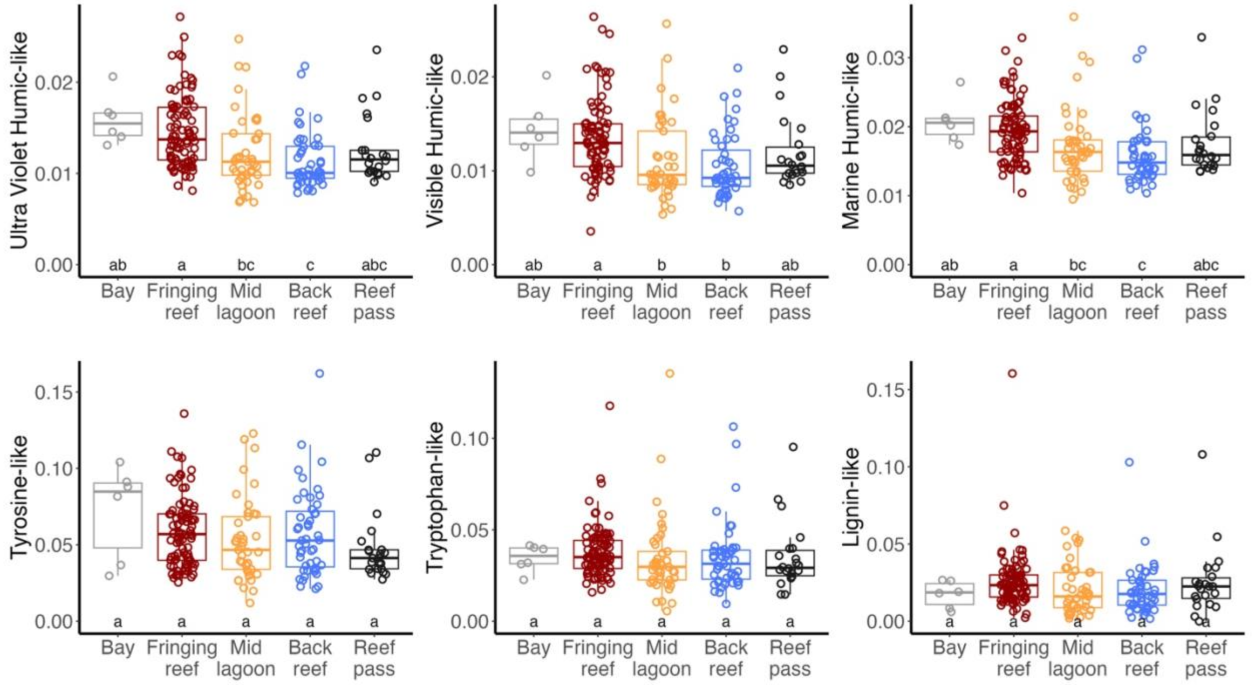


| *Parameter* | *term* | *Df* | *Sum.Sq* | *Mean.Sq* | *F* | *p* | |
|---|---|---|---|---|---|---|---|
| *Ultra-violet humic-like* | Habitat | 4 | 0.00127 | 0.00032 | 10.163 | 0 | *** |
| | Year | 1 | 0.00154 | 0.00154 | 49.403 | 0 | *** |
| | Habitat:Year | 4 | 0.00013 | 3e-05 | 1.045 | 0.383 | |
| | Residuals | 552 | 0.01721 | 3e-05 | | | |
| *Visible humic-like* | Habitat | 4 | 0.00111 | 0.00028 | 7.901 | 0 | *** |
| | Year | 1 | 0.0031 | 0.0031 | 88.087 | 0 | *** |
| | Habitat:Year | 4 | 0.00011 | 3e-05 | 0.763 | 0.55 | |
| | Residuals | 552 | 0.0194 | 4e-05 | | | |
| *Marine humic-like* | Habitat | 4 | 0.00145 | 0.00036 | 7.232 | 0 | *** |
| | Year | 1 | 0.00741 | 0.00741 | 147.64 | 0 | *** |
| | Habitat:Year | 4 | 0.00039 | 1e-04 | 1.925 | 0.105 | |
| | Residuals | 552 | 0.02771 | 5e-05 | | | |
| *Tyrosine-like* | Habitat | 4 | 0.0088 | 0.0022 | 1.3 | 0.269 | |
| | Year | 1 | 0.23493 | 0.23493 | 138.771 | 0 | *** |
| | Habitat:Year | 4 | 0.01309 | 0.00327 | 1.933 | 0.103 | |
| | Residuals | 552 | 0.93451 | 0.00169 | | | |
| *Tryptophan-like* | Habitat | 4 | 0.00221 | 0.00055 | 0.691 | 0.598 | |
| | Year | 1 | 0.01798 | 0.01798 | 22.463 | 0 | *** |
| | Habitat:Year | 4 | 0.00465 | 0.00116 | 1.452 | 0.216 | |
| | Residuals | 552 | 0.44177 | 8e-04 | | | |
| *Lignin-like* | Habitat | 4 | 0.00614 | 0.00154 | 1.62 | 0.168 | |
| | Year | 1 | 0.01657 | 0.01657 | 17.483 | 0 | *** |
| | Habitat:Year | 4 | 0.00352 | 0.00088 | 0.928 | 0.447 | |
| | Residuals | 552 | 0.52322 | 0.00095 | | | |

Supporting information for:
Terrigenous inputs link nutrient dynamics to microbial communities in a tropical lagoon

**fDOM indices**

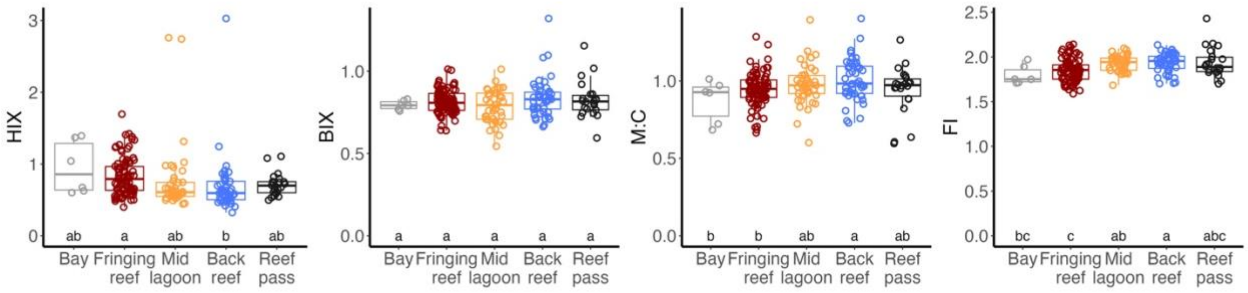


| *Parameter* | *term* | *Df* | *Sum.Sq* | *Mean.Sq* | *F* | *p* | |
|---|---|---|---|---|---|---|---|
| *HIX* | Habitat | 4 | 3.04424 | 0.76106 | 2.806 | 0.025 | * |
| | Year | 1 | 3.3693 | 3.3693 | 12.424 | 0 | *** |
| | Habitat:Year | 4 | 0.21429 | 0.05357 | 0.198 | 0.94 | |
| | Residuals | 551 | 149.4221 | 0.27118 | | | |
| *BIX* | Habitat | 4 | 0.14009 | 0.03502 | 1.55 | 0.186 | |
| | Year | 1 | 1.16464 | 1.16464 | 51.554 | 0 | *** |
| | Habitat:Year | 4 | 0.14829 | 0.03707 | 1.641 | 0.162 | |
| | Residuals | 552 | 12.46999 | 0.02259 | | | |
| *M:C* | Habitat | 4 | 0.45444 | 0.11361 | 1.648 | 0.161 | |
| | Year | 1 | 70.68102 | 70.68102 | 1025.367 | 0 | *** |
| | Habitat:Year | 4 | 1.24471 | 0.31118 | 4.514 | 0.001 | *** |
| | Residuals | 552 | 38.0507 | 0.06893 | | | |
| *FI* | Habitat | 4 | 0.98176 | 0.24544 | 6.573 | 0 | *** |
| | Year | 1 | 12.97473 | 12.97473 | 347.461 | 0 | *** |
| | Habitat:Year | 4 | 0.14332 | 0.03583 | 0.96 | 0.429 | |
| | Residuals | 552 | 20.61252 | 0.03734 | | | |

Supporting information for:
Terrigenous inputs link nutrient dynamics to microbial communities in a tropical lagoon

**Supplementary materials S10. Within-year water chemistry ordination**
Water chemistry ordinations for all sampling within individual years show consistent variability among samples across years.

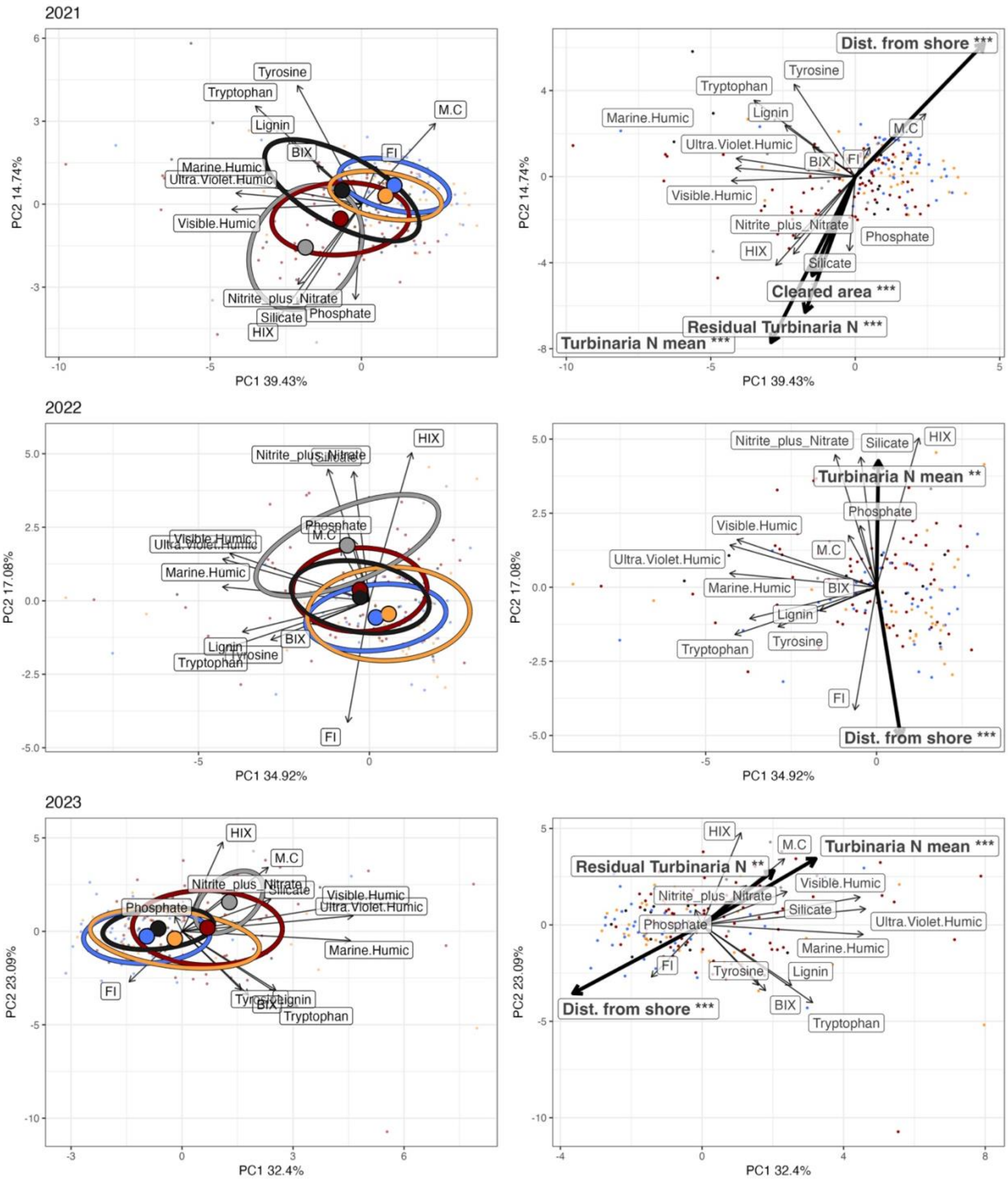

Supporting information for:
Terrigenous inputs link nutrient dynamics to microbial communities in a tropical lagoon

**Supplementary materials S11. Within-year microbial community ordinations**
Microbial community ordinations for each year show consistent variability in community structure by habitat (a; annotated letters indicate differences in dispersion among habitats) and associations with environmental variables (b) across the three years of the study.

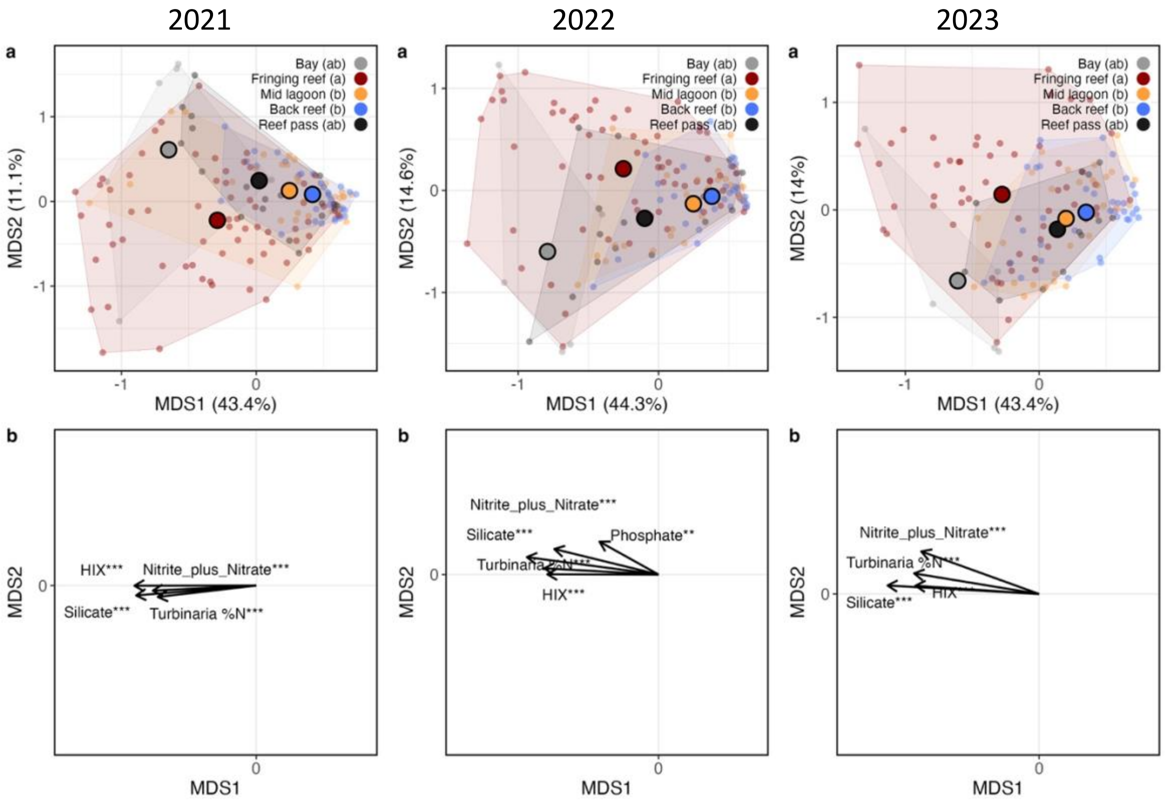

Supporting information for:
Terrigenous inputs link nutrient dynamics to microbial communities in a tropical lagoon
